# MicroRNA miR-7 regulates synthesis and secretion of insulin-like peptides

**DOI:** 10.1101/791640

**Authors:** Pamela Agbu, Justin J. Cassidy, Jonathan Braverman, Alec Jacobson, Richard W Carthew

## Abstract

The insulin/IGF pathway is essential for linking nutritional status to growth and metabolism. MicroRNAs (miRNAs) are short RNAs that are players in the regulation of this process. The miRNA miR-7 shows highly conserved expression in insulin-producing cells across the animal kingdom, however, its conserved functions in regulation of insulin-like peptides (ILPs) remains unknown. Using *Drosophila* as a model, we demonstrate that miR-7 limits ILP availability by inhibiting its production and secretion. Increasing miR-7 alters body growth and metabolism in an ILP-dependent manner, elevating circulating sugars and total body triglycerides, while decreasing animal growth. These effects are not due to direct targeting of ILP mRNA, but instead arise through alternate targets that affect the function of ILP-producing cells. The *Drosophila* F-actin capping protein, CPA, is a direct target of miR-7, and knockdown of CPA in IPCs phenocopies the effects of miR-7 on ILP secretion. This regulation of CPA is conserved in mammals, with the mouse ortholog Capza1 also targeted by miR-7 in β-islet cells. Taken together, these results support a role for miR-7 regulation of an actin capping protein in insulin regulation, and highlight a conserved mechanism of action for an evolutionarily ancient microRNA.

**Disclosure Statement:** PA, JB, AJ and RWC have nothing to declare. JJC is an employee of Sg2, LLC, a healthcare consulting company.

## Introduction

Adult diabetes is a metabolic disease characterized by defects in insulin production and utilization. Millions of people are diagnosed with Type II diabetes worldwide, and the numbers of those afflicted are predicted to steadily increase^1^. Progression of Type II diabetes involves the interplay between insulin and blood glucose levels, with initial disease stages characterized by insulin resistance and hyperinsulinemia^2^. In more advanced stages of Type II diabetes, insulin secretory ability becomes further compromised and results in hyperglycemia^3^. The net result is a dysfunction in carbohydrate homeostasis.

Insulin regulates glucose metabolism in the classical insulin-responsive tissues such as adipose, muscle and liver^4^. However, insulin and related peptides, called insulin-like growth factors (IGFs), also regulate growth and body size during the embryonic, fetal, and juvenile phases of life^5–7^. Growth depends upon nutrient availability, and insulin acts to maintain carbohydrate consumption by cells for their continued growth and proliferation. IGFs stimulate cell growth by activating the PI3K-TOR transduction pathway, and thereby increase protein synthesis capacity within cells^8^.

The fruit fly *Drosophila melanogaster* has proven to be a useful model to study questions related to insulin/IGF effects on growth and metabolism^9,10^. One function of the insulin/IGF and TOR systems in *Drosophila* is to assess the nutritional status of the growing organism and to relay this information into developmental decisions that appropriately adjust the growth of the organism to its final specified size. Growth is facilitated by the *Drosophila* insulin-like peptides (dILPs), with dILP2 being the most potent regulator of growth. Other dILPs, including dILP3 and dILP5, have also been shown to promote growth^10,11^. dILP2 is most highly related to insulin, with 35% sequence identity, and dILP2 is sufficient to restore normal circulating sugar levels in diabetic flies^9,12^. dILP5 is slightly less similar to insulin, but structural analysis has demonstrated that dILP5 shares the basic fold of the insulin peptide family^13^. Moreover, dILP5 binds to the mammalian Insulin Receptor and can lower blood glucose in rats^13^. The dILPs bind a single *Drosophila* insulin receptor to activate the highly conserved MAPK and TOR pathways^12,14^. Inhibiting these pathways or genetic ablation of *dILP* genes result in decreased growth rate, developmental delay, and metabolic abnormalities resembling a diabetic-like condition^11^. Changes in mutant animal growth rate during larval life result in adults with altered body size^9^.

The primary site of insulin synthesis in mammals is the β-cells of the pancreas. The transcription factors PDX-1, MaFA, Beta2/NeuroD1, and Pax6 function in mature β-cells to activate insulin gene expression in response to glucose-stimulated activation of the MAPK pathway^15^. Although gene expression affects insulin peptide levels, it is primarily insulin release from β-cells by exocytosis that dictates the level of circulating insulin^16^. Glucose metabolism in β-cells increases the ATP:ADP ratio, activating ATP-sensitive potassium channels that trigger plasma membrane depolarization and calcium influx^17–20^. This leads to a number of downstream events, including fusion of insulin-containing secretory granules with the membrane, and subsequent release of insulin into circulation^17^.

Although *Drosophila* does not have a pancreas, flies have a group of 14 secretory neurons located in the brain, known as insulin producing cells (IPCs). These secrete dILP2, dILP3 and dILP5^10^. Expression and release of IPC dILPs are nutritionally regulated via signals from the fat body, glia, and corpora cardiaca^21–23^. These signals form a complex and dynamic regulatory network that converges on the IPCs and coordinates growth with nutritional status. Some of the transcription factors important for mammalian β-cell function are also important for fly IPCs. Mutation of the *Drosophila Pax6* ortholog in IPCs results in decreased transcription of *dILP5*, similar to what is seen in mammals with Pax6 and insulin gene transcription^24^. The insulin secretory mechanism has also been conserved in *Drosophila*. Nutrient sensing leads to downstream events such as potassium channel closure and calcium influx prior to dILP release^21,25,26^.

MicroRNAs (miRNAs) are small non-coding RNAs that are transcribed from genes and processed from long precursor RNAs into 22-nucleotide single strands. They form a miRISC complex with Argonaute proteins, and guide the complex to associate with target mRNAs, destabilize them, and reduce their translation capacity^27^. As such, miRNAs serve as weak repressors of gene expression. There are hundreds to thousands of distinct miRNAs encoded by animal genomes, and they regulate a majority of the protein-coding transcriptome^27^. MicroRNAs have been found to control insulin production and secretion in mammals^28–30^. Ablation of Dicer, an enzyme necessary for the biogenesis of most miRNAs, results in defective insulin production and secretion in mice^31^. Evidence also exists that individual miRNAs play a role. The mouse miRNA miR-7 inhibits glucose-stimulated insulin secretion through regulation of SNCA, a promoter of SNARE oligomerization^28^. miR-7 expression is induced in β-cells by Ngn3 and NeuroD/Beta2^32^.

miR-7 is an evolutionarily ancient miRNA, having arisen in the ancestor of all bilaterian animals^33^. The sequence of miR-7 is perfectly conserved between *Drosophila melanogaster* and various mammalian species, suggesting strong functional conservation (Fig. 1A). Moreover, miR-7 shows conserved expression in neurosecretory cells of invertebrates and vertebrates^34^. However, it remains unclear whether miR-7’s role in insulin function is similarly conserved in *Drosophila*. Here, we demonstrate that miR-7 is a key regulator of *Drosophila* dILP biology, and like many of the transcription factors functioning in pancreatic β-cells, miR-7 exerts its functions at multiple levels, acting in both the production and in the secretory processes. We identify the gene encoding F-actin capping protein alpha, CPA, as a direct target of miR-7 repression. CPA appears to regulate dILP2 release from IPCs thereby affecting circulating levels of dILP2. Moreover, the mouse homolog of CPA, CAPZA1, is a miR-7 target in pancreatic β-cells, suggesting that miR-7 employs a conserved mechanism of action in regulating insulin secretion across the animal kingdom.

**Figure 1.**
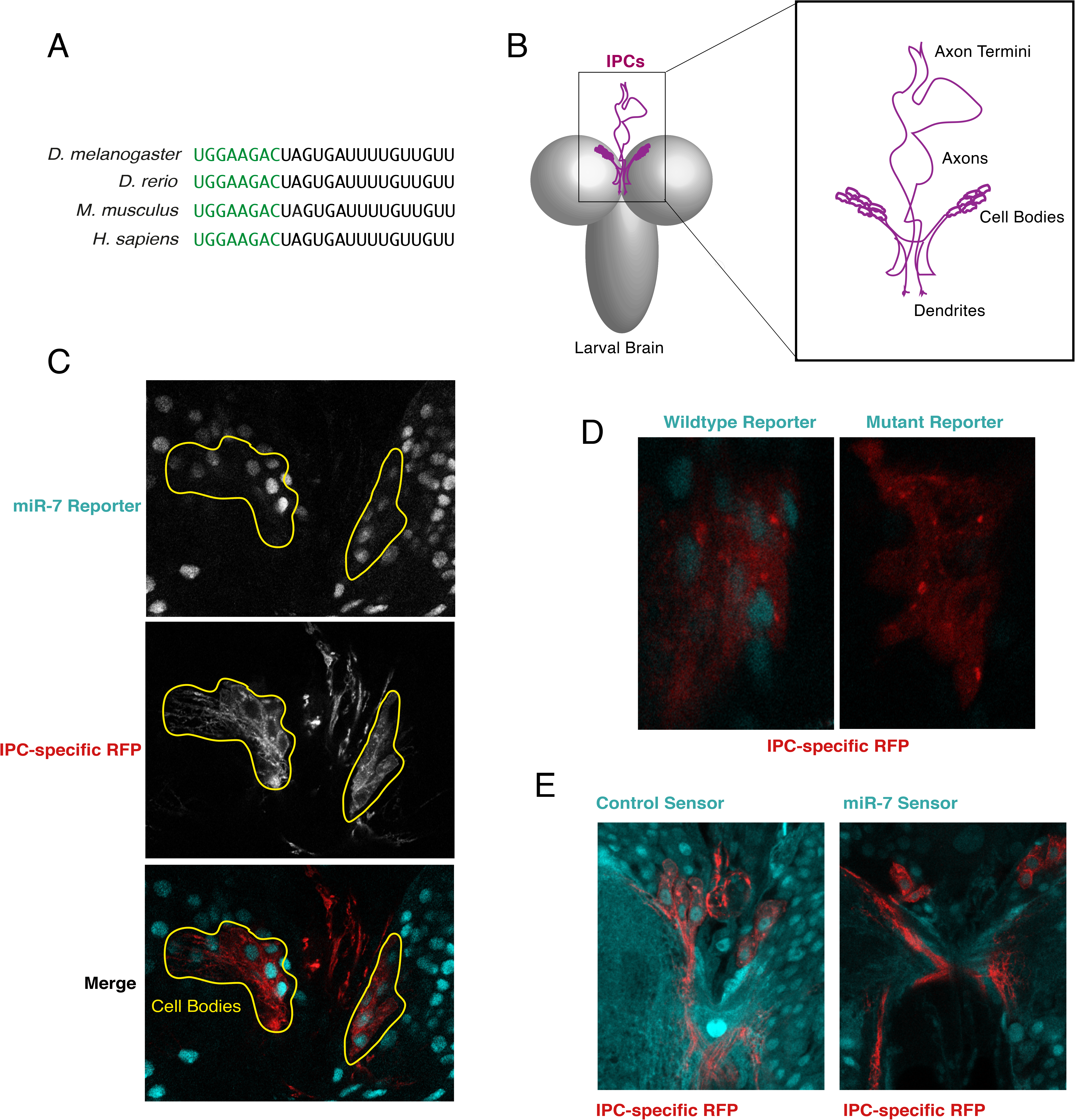
miR-7 is expressed and active in IPCs. (**A**) Alignment of the mature miR-7-5p sequence (5’ - 3’) from different animal species. Highlighted in green is the seed sequence. Shown are the miR-7a-1 (zebrafish and mouse) and miR-7-1 (human) paralogs. (**B**) Schematic of the location and morphology of the two clusters of IPCs in the larval brain. Each cluster contains seven cells. Anterior is top and dorsal is face up. (**C**) Transgene reporter of *mir-7* enhancer activity as reported by nuclear GFP. Two clusters of IPC cell bodies are specifically highlighted by RFP fluorescence in the larval brain. (**D**) An IPC cluster expressing the GFP reporter that either contains a wildtype *mir-7* enhancer or a mutant enhancer in which two bHLH binding sites are altered. (**E**) Transgenic sensors expressing GFP under constitutive promoter control. Right, a sensor containing two perfect binding sites for miR-7 in its 3’UTR. Left, a sensor lacking such binding sites. Each image shows two IPC clusters from a larval brain, marked by RFP fluorescence. For panels C - E, anterior is top.

## Methods

### Genetics

For all experiments, *Drosophila melanogaster* was raised using standard lab conditions and food. Stocks were either obtained from the Bloomington Stock Center, from listed labs, or were derived in our laboratory. To generate *mir-7* null mutant animals, we crossed two different deletion mutants together and made a trans-heterozygous *mir-7*^Δ1^/*Df(2R)exu1* mutant. To specifically misexpress transgenes in IPCs, we used *dILP2-Gal4* transgenic lines. *dILP2-Gal4* fuses the *insulin-like peptide 2* gene promoter to Gal4, and specifically drives its expression in brain IPCs^9^. We used either a *dILP2-Gal4* line with a transgenic insertion on chromosome II or III. To genetically ablate the IPCs of the brain, *w*^1118^ animals were constructed bearing a *dILP2-Gal4* gene on chromosome III and a *UAS-Reaper* (*Rpr*) gene on chromosome II. *Rpr* is a pro-apoptotic gene that is sufficient to kill cells in which it is expressed^35^. Examination of *dILP2-Gal4 UAS-Rpr* larval brains showed that they almost completely lacked IPCs (data not shown).

To overexpress miR-7 in IPCs, we used a *UAS-mir-7* transgene in which a 432 bp DNA fragment containing the miR-7 precursor is downstream of the UAS-driven promoter^36^. This was combined with the *dILP2-Gal4* transgene located on chromosome III. To knockdown miR-7 in IPCs, we used a *UAS-mir-7 sponge* transgene in which the UAS-driven promoter sits upstream of GFP coding sequence and a sponge sequence in the 3’UTR^37^. The sponge sequence consists of 10 tandem binding sites for miR-7, with base mismatches at positions 9-12. This *UAS-mir-7 sponge* transgene was combined with the *dILP2-Gal4* transgene located on chromosome II. For all wildtype controls involving IPC-specific expression of transgenes, we used animals that contained the matching *dILP2-Gal4* transgenic line.

To mark IPCs, we used either *UAS-mCD8::mRFP* or *UAS-DsRed.nls*. The former expresses a mouse CD8 transmembrane domain fused to the N-terminus of mRFP^38^. The latter expresses dsRed with a nuclear localization tag at the C-terminus^39^.

To determine whether miR-7 RNA is expressed in IPCs, we used a *mir-7* transcriptional enhancer driving GFP^40^. This consists of a 349 bp region upstream of the *mir-7* transcription start site. The *mir-7* enhancer region was placed upstream of a minimal promoter (TATA) and sequence encoding for a nuclear localized GFP was placed downstream of this region. The *mir-7* enhancer contains two binding sites for the bHLH factor Atonal, which was previously demonstrated to regulate enhancer activity in the eye^40^. To determine if bHLH factors regulate of enhancer activity in IPCs, we used animals that expressed GFP under the control of the *mir-7* enhancer, but with the two bHLH binding sites mutated^40^.

To assay miR-7 silencing activity in IPCs, we utilized miRNA silencing sensors. The miR-7 sensor expresses GFP under the control of the tubulin promoter and with a 3’UTR that contains two perfectly complementary miR-7 binding sites^41^. The control sensor drives GFP under the control of the same tubulin promoter, however there are no miR-7 binding sites in the 3’UTR^41^.

To quantify stored and circulating dILP2 levels, we used flies expressing dILP2HF^25^. These flies are null mutant for the endogenous *dILP2* gene, but express a genomic *dILP2* transgene that has been tagged with an HA epitope at the B-chain carboxy terminus and a FLAG epitope at the A-chain amino terminus.

To identify miR-7 regulatory targets, RNAi lines were obtained from Bloomington Drosophila Stock Resource Center or from the Vienna Drosophila Resource Center. Each RNAi line has a UAS promoter that drives a double-stranded hairpin RNA corresponding to an annotated *Drosophila* gene^42^. Each line was crossed to *dILP2-Gal4* animals, and progeny bearing both transgenes were assayed.

## Body Weight Measurement

All *Drosophila* to be weighed were raised at 25°C. Adult females were weighed no more than 24 hours after eclosion, and batches of 20-50 animals were counted and weighed on a Mettler analytical balance. Larvae undergoing the pupal molt were identified as "white prepupae", and batches of these animals were counted and weighed as described. Mean weight was estimated for each replicate batch, and multiple replicates were assayed for both adult and larval experiments. For all weighing experiments, wildtype controls were animals with one copy of the *dILP2-Gal4* gene located on either the second or third chromosome. The insertion site of the *dILP2-Gal4* gene in control lines was matched to its *mir-7* perturbation since we found that different insertion lines caused animals to vary in weight. The cause might be linked to Gal4 itself mildly affecting IPC function

### Adult Wing Measurement

Length of the adult wing scales proportionally with total size of adult *Drosophila*. To measure wing length, wings from 3-5 day old adult male flies were preserved in 70% (v/v) ethanol. Preserved wings were mounted in 70% (v/v) glycerol and imaged using a Zeiss Axiophot microscope and a 4x objective. Images were digitized using the Zeiss Zen imaging software. To measure wing blade length, the distance between the anterior crossvein and wing margin (point where it intersects with the L3 longitudinal vein) was measured using the Microruler package in ImageJ software.

### Metabolite Measurements

#### Triglycerides

To determine total body triglycerides, five wandering third instar larvae were homogenized in freshly made phosphate buffered saline (PBS) + 0.05% (v/v) Tween-20 (Sigma).. Homogenate was heated to 70°C, centrifuged at 3000 x g, and then at 3500 x g. Following centrifugation, 25 μl supernatant was added to 100 μl Infinity Triglycerides Reagent (Thermo Scientific) and incubated at 37°C for 10 minutes (min). Absorbance at 540 nm was measured to determine total triglycerides. Following triglyceride measurements, total protein was determined by the Bradford assay.

To determine levels of circulating triglycerides, hemolymph was extracted from wandering third instar larvae as described^43^ and diluted 1:10 in PBS. Samples were heated at 70°C for 5 min and then centrifuged for 10 min at 16000 x g. Five μl supernatant was added to 100 ul Infinity triglyceride reagent, and samples were incubated at 37°C for 10min. Absorbance at 540 nm was measured using a Biotek Synergy 4 microplate reader.

#### Glucose

To measure circulating glucose levels, porcine kidney trehalase (Sigma) was diluted 1:1000 (v/v) in Infinity Glucose Reagent (Thermo Scientific), and pH was adjusted to 6.5 with HCl. One μl of hemolymph was extracted from wandering third instar larvae and added to 100 μl Glucose Reagent supplemented with trehalase, and samples were incubated overnight at 37°C. Absorbance at 340 nm was measured using the Biotek Synergy4 microplate reader.

### ELISA assay of brain and circulating dILP2HF

To measure circulating dILP2HF levels, 2.5 μg/ml anti-FLAG antibody (Sigma-Aldrich F1804) in 0.2 M sodium bicarbonate pH 9.4 buffer was incubated overnight in wells of an ELISA plate (Nunc Immuno Module, Thermo Scientific 468667). Following two washing steps, wells were blocked overnight in PBS + 2% (w/v) BSA (Sigma). Three μl of hemolymph was extracted from male wandering third instar larvae and added to 37 μl PBS. Hemolymph samples were centrifuged at 16000 x g for 5 min. 35 μl of supernatant was added to 5 μl PBS + 2% (v/v) Tween-20 supplemented with anti-HA-antibody conjugated to horseradish peroxidase (Roche) (1:500 v/v dilution from stock). This was added to one well of a blocked ELISA plate, and incubated overnight at 4°C. Wells were washed with PBS+ 0.2% (v/v) Tween-20 and developed using 100 μl One Step Ultra TMB – ELISA Substrate (Thermo Scientific) for 30 minutes at room temperature. Reactions were stopped with 100 μl 2 M sulfuric acid, and absorbance was measured at 450 nm.

To measure stored brain dILP2HF levels, three brains were dissected from wandering third instar larvae and homogenized in 150 μl PBS + 1% (v/v) Triton-X100. Samples were vortexed for 5 min and then centrifuged at 16000 x g for 5 min. 35 μl supernatant was used to perform the ELISA assay, as described above. Total brain protein levels were determined by Bradford assay.

### Tissue preparation and imaging

Brains of male wandering third instar larvae were dissected and fixed for 60 min at room temperature in 4% (w/v) paraformaldehye (Sigma) in PBS. Washes were performed using PBS. Brain tissue was mounted in Vectashield and images were collected using a Leica SP5 laser scanning confocal microscope. Fluorophores were excited sequentially using the 488 and 561 laser lines. When comparing conditions, samples were imaged side by side on the same slide, using identical parameters between conditions. Images were collected as Z-stacks, with optical section thickness set to 2.98 μm for miR-7 sensor and reporter images, while images of the miR-7 sponge had an optical section thickness of 2.52 μm. Imaging was performed with the 40X oil objective lens, with 1.25 NA, and 1024 x 1024 resolution. All scans were done with the bidirectional scanner, and line averaging was set to 6x. Offset was set to zero, and gain was adjusted to minimize overexposure of the brightest sections.

### RNA extraction and RT-qPCR

Seven to ten brains from male wandering third instar larvae were dissected in PBS and homogenized in 200 μl TRIzol Reagent (Invitrogen). RNA was extracted according to the manufacturer’s instructions. SuperScript III reverse transcriptase (Thermo) was used for reverse transcription of RNA. The reaction was primed using oligo-dT and random 9mer primers (2:1 molar ratio). 25 ng cDNA was used as a template for each qPCR reaction using SyBr Green. Gene mRNA levels were normalized to β-tubulin mRNA level using the standard delta-delta Ct method.

### Prediction of miR-7 targets

The algorithms PicTar, Sloan-Kettering, PITA, MiRTE, miRanda, and TargetScan were used to predict miR-7 targets in annotated *Drosophila melanogaster* genes^40^. For prediction of miR-7 targets in annotated human genes, PicTar, TargetScan, and PITA were used^40^. Genes predicted by three or more algorithms for *Drosophila*, and two or more algorithms for humans were considered to be predicted targets. Human orthologs of *Drosophila* predicted targets were determined using Ensembl, InParanoid, and Orthology Matrix^40^.

### Luciferase Assays of miR-7 sites in the CPA 3’UTR

The annotated 3’UTR of the *cpa* gene (2R:21059062..21059883 [+], Dmel release 5.57) was synthesized as a gBlock (IDT) and inserted into the pMT-GL3 plasmid, which expresses the firefly luciferase gene under the control of a copper-inducible metallothionein promoter^44^. Construction of this reporter involved swapping the *Drosophila cpa* 3’UTR for the SV40 3’UTR present in pMT-GL3. Each predicted miR-7 binding site in the *cpa* 3’UTR was mutated singly or together to generate three different mutant reporters. Site 1 (2R:21059118-21059126) was mutated from GTCTTCCA to GCTCCCCA. Site 2 (2R:21059595-21059601) was mutated from GTCTTT to GTGGGG.

To perform luciferase assays, *Drosophila* S2 cells were cultured in Schneider’s media supplemented with 10% FBS and seeded at a density of 2×10^6^ cells per ml prior to transfection. Cells were co-transfected with pMT-Ren, which is a control reporter for Renilla luciferase^44^, and pMT-GL3 containing either a wildtype or mutant *cpa* 3’UTR. These were transfected at a 1:10 molar ratio, respectively, Cells were also transfected with pUAS-miR-7 and Ubiquitin-Gal4 plasmids. Transfections were performed with Effectene Transfection Reagent (Qiagen) according to manufacturer’s protocol. To induce reporter expression, Cu_2_SO_4_ was added to 0.5 mM final concentration. Twenty four hours after induction, cells were lysed, and luciferase assays were performed using the Dual Luciferase Assay system (Promega). Luciferase activities were quantified using the Glomax Luminometer, and Firefly luciferase activity was normalized as the ratio of Firefly to Renilla enzymatic activities per sample.

### Analysis of Mouse β-cell Transcriptomics

Microarray data was downloaded from GEO (accession no. GSE48195)^28^. Scatterplots were rendered in Prism 8 (Graphpad).

## Results

miR-7 is expressed in insulin-producing cells of vertebrate species^28,45,46^ However, it is unclear whether miR-7 is expressed in the IPCs of *Drosophila*. In order to answer this question, we examined a transgenic reporter gene for miR-7 expression. This transgene has the transcriptional enhancer of the *mir-7* gene fused to a minimal promoter driving transcription of GFP. The reporter faithfully reproduces the expression pattern of miR-7 in various tissues of the fly^40^. The transgene showed broad expression in the larval brain, as evident by nuclear-localized GFP fluorescence. To mark the location of the IPCs, we expressed membrane-bound RFP specifically in IPCs using the *dILP2* gene promoter to drive expression. The IPCs occupy a stereotyped position in the larval brain with 14 cells residing on the left and right hemispheres of the brain (Fig. 1B). The IPCs could clearly be identified with the membrane RFP marker (Fig. 1C), and all IPCs were labeled with nuclear GFP, suggesting that the *mir-7* enhancer is active in IPCs (Fig. 1C). To further validate our conclusion, we examined GFP expression from a reporter gene in which two binding sites for bHLH transcription factors were mutated in the enhancer. These binding sites are essential for *mir-7* expression in other tissues of *Drosophila*^40^. The mutant enhancer was completely inactive in the larval IPCs (Fig. 1D). Thus, the *Drosophila mir-7* gene requires bHLH factors to augment *mir-7* expression in IPCs. This is comparable to the requirement of the bHLH factor NeuroD/Beta2 for *mir-7* transcription in mouse β-cells^32^.

To demonstrate that miR-7 RNA is functionally active in IPCs, we used a transgenic sensor for miR-7 silencing activity^41^. Transgenic mRNA is transcribed ubiquitously in all cells, and it contains two perfect binding sites for miR-7. If miR-7 is loaded into miRISC, it inhibits the synthesis of the GFP protein product of the transgene in that cell. Thus, reduction of GFP fluorescence is an indicator of miR-7 silencing activity. The miR-7 sensor showed weak GFP expression in larval IPCs (Fig. 1E). As a control, a transgene sensor lacking miR-7 binding sites was also examined for GFP expression in IPCs. As expected, the control sensor gave strong ubiquitous GFP expression in larval IPCs (Fig. 1E). Thus, miR-7 is not only expressed but is functionally active in the IPCs of growing *Drosophila*.

### IPC-specific miR-7 regulates body weight and metabolite levels

Growth in *Drosophila* is facilitated by the IPCs in the larval brain. Ablation of larval IPCs results in smaller adults^9^. IPC ablation can be engineered by expressing the pro-apoptotic gene *Reaper* in IPCs using *dILP2-Gal4*. We wondered if miR-7 plays a role in IPC regulation of growth. Therefore, we made gain-of-function and loss-of-function perturbations of miR-7 specific to IPCs and not to other cells of the body. We reasoned that since miR-7 is expressed in many tissues^40^, classical *mir-7* mutations might produce confounding effects not directly related to IPC function. We overexpressed miR-7 specifically in IPCs by combining *dILP2-Gal4* and *UAS-mir-7* transgenes in animals. Since Gal4 is only present in IPCs due to the *dILP2* gene promoter, miR-7 is only overexpressed in these cells. We weighed such animals when they reached adulthood, and observed a decrease in mean body weight compared to wildtype siblings (Fig. 2A). We compared the effects of miR-7 overexpression with IPC cell ablation by *Reaper*. The effect of miR-7 overexpression on adult body weight was slightly less potent than the effect of IPC ablation on body weight (Fig. 2A).

**Figure 2.**
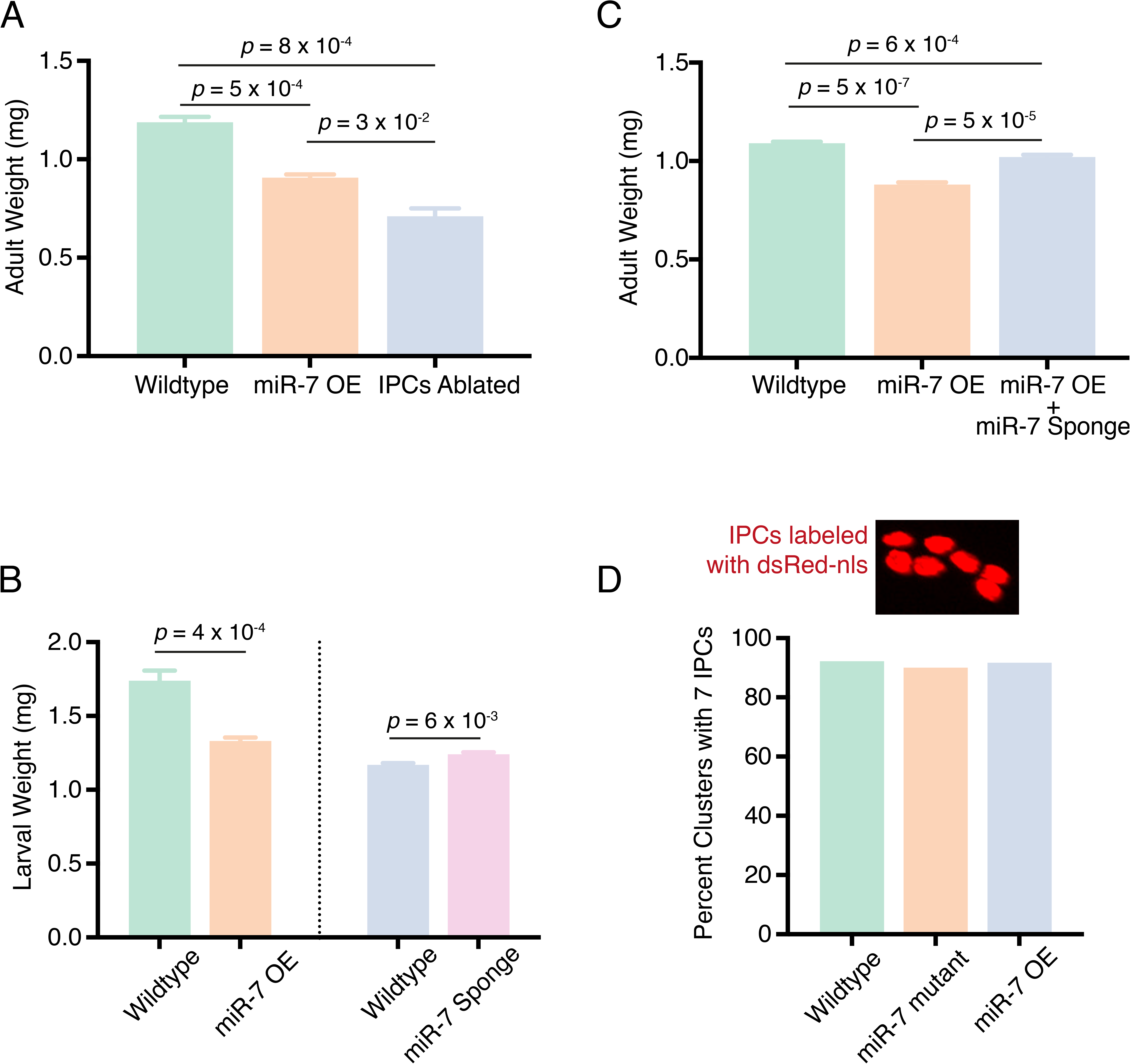
mir-7 mutants alter body weight. (**A**) Mean adult weight of females that have miR-7 overexpression (OE) in IPCs or have had their IPCs ablated. (**B**) Mean weight of female larvae at the pupal molt that have miR-7 overexpression (OE) or the miR-7 Sponge expressed in IPCs. Each condition is paired with a wildtype control line that contains the *dILP2-Gal4* driver used in each experiment. Different *dILP2-Gal4* drivers affect body weight on their own, which requires rigorous pairing for controls. (**C**) Mean adult weight of females that have miR-7 overexpression (OE) compared to wildtype. One set of miR-7 OE animals also had the miR-7 Sponge co-expressed in their IPCs. (**D**) Shown on top is nuclear dsRed fluorescence in IPCs within one larval brain cluster. Note there are seven nuclei. At bottom is the percent of larval brain clusters having seven IPCs from animals with different genotypes. Number of scored clusters range from 12 to 64. A Chi-Square test on the data showed that none of the treatments had significantly different percentages (*p* = 0.93). For panels A - C, error bars represent the standard error of the mean. *P* values are derived from unpaired two-tailed student T-tests.

IPCs primarily regulate adult body size by affecting the growth rate during the juvenile phase of the life cycle ^9^. To determine whether miR-7 regulates larval growth, we weighed animals at the larval-to-pupal molt. As observed in adults, miR-7 overexpression caused a decrease in body size at this stage (Fig. 2B).

To generate a loss-of-function perturbation, we inhibited miR-7 by overexpressing a miR-7 sponge RNA specifically in IPCs. Sponge RNAs contain partially complementary binding sites to a miRNA of interest, and are produced from transgenes within cells^37^. When overexpressed, sponge RNAs sequester the miRNA of interest and titrate it away from its natural targets. The *UAS-miR-7 sponge* transgene expresses mRNA with ten tandem copies of a sequence complementary to miR-7 inserted in the 3’ UTR^37^. To confirm the potency of the miR-7 sponge, we co-expressed the sponge along with *UAS-mir-7* in IPCs using *dILP2-Gal4*. This largely neutralized the effect of miR-7 overexpression on body weight (Fig. 2C). When we expressed just the miR-7 sponge alone in IPCs, there was a small but significant increase in body weight of animals at the larval-to-pupal molt (Fig. 2B). Thus, both loss-of-function and gain-of-function experiments indicate that IPC-specific miR-7 inhibits growth of *Drosophila*.

One possible role for miR-7 in IPCs might be for their development and survival. If so, we predicted that *mir-7* mutants would display an abnormal number or morphology of IPCs. However, null *mir-7* mutants and *mir-7* overexpression mutants had the normal number and arrangement of IPCs (Fig. 2D).

Growth is controlled by the balanced utilization and storage of carbohydrates and lipids. Impaired insulin signaling in diabetic patients is characterized by fasting hyperglycemia. Late third instar *Drosophila* larvae undergo cycles of feeding and fasting as they prepare for the pupal molt. Fasting animals can be easily identified by their clear digestive systems when given colored food. We extracted hemolymph from fasting late third instar larvae and measured their circulating sugar levels. Insects have two types of carbohydrates in circulation: glucose and trehalose. Glucose is obtained from food sources, while trehalose (the disaccharide of glucose) originates from the fat body, and is the primary sugar consumed by cells. Although knockdown of miR-7 in IPCs did not have a significant effect on circulating sugar levels, overexpression of miR-7 in IPCs caused an increase in circulating sugars (Fig. 3A). Inhibition of dILP expression or ablation of IPCs generate similar effects on circulating sugars^9,11^.

**Figure 3.**
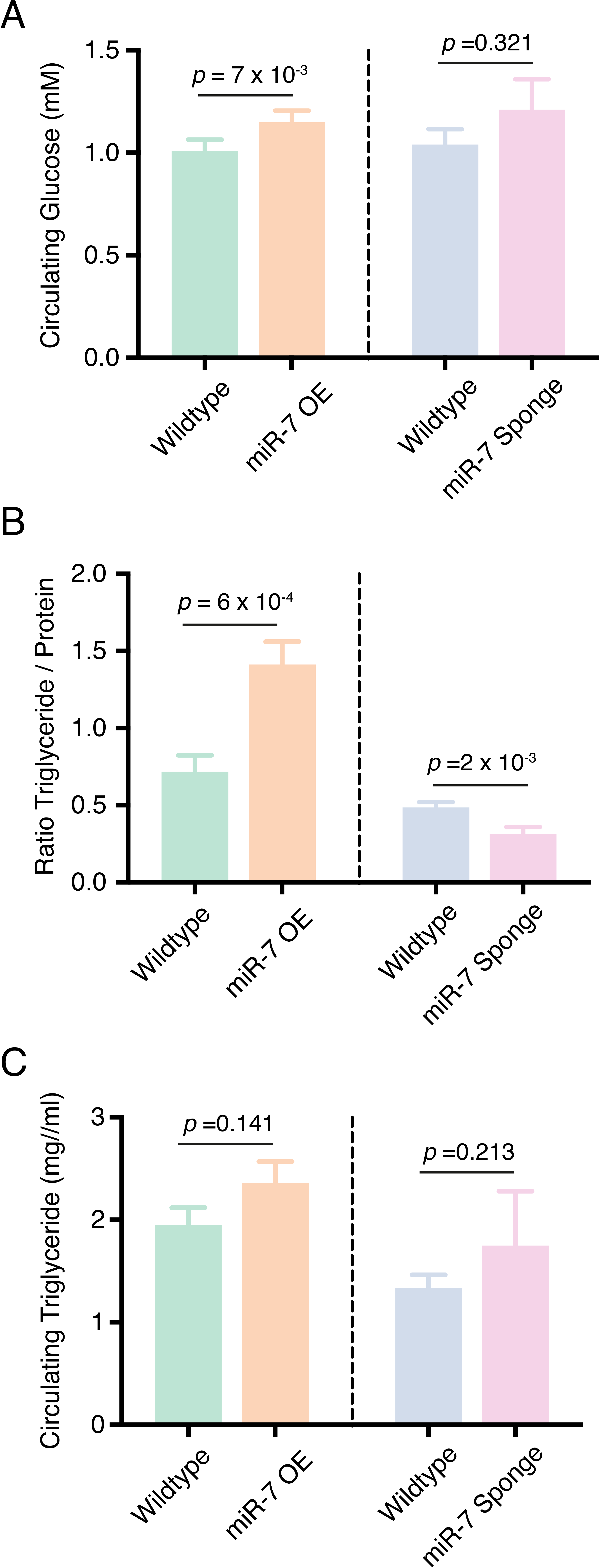
mir-7 mutants alter carbohydrate and lipid metabolism. (**A**) Mean concentration of glucose (monosaccharide plus trehelose disaccharide) in larval hemolymph from animals that have miR-7 overexpression (OE) or the miR-7 Sponge expressed in IPCs. Each condition is paired with a wildtype control line that contains the *dILP2-Gal4* driver used in each experiment. (**B**) Average ratio of total body triglyceride to protein (w/w) in larvae that have miR-7 overexpression (OE) or the miR-7 Sponge expressed in IPCs. Each condition is paired with a wildtype control line that contains the *dILP2-Gal4* driver used in each experiment. (**C**) Mean concentration of triglyceride in larval hemolymph from animals that have miR-7 overexpression (OE) or the miR-7 Sponge expressed in IPCs. Each condition is paired with a wildtype control line that contains the *dILP2-Gal4* driver used in each experiment. Error bars represent the standard error of the mean. *P* values are derived from unpaired two-tailed student T-tests.

The storage of fat in insects is primarily in the form of triglycerides. The ratio of body triglyceride to protein is a measure of relative fat storage in *Drosophila*. We compared this ratio between miR-7 overexpressing and wildtype larvae. There was a significant increase in normalized triglycerides when miR-7 was overexpressed in IPCs (Fig. 3B). This phenotype is also observed when adults lack any IPCs^47^. Conversely, expression of miR-7 sponge RNA in IPCs caused a decrease in total triglycerides (Fig. 3B). These effects were exerted on non-circulating triglycerides since there was little or no difference in circulating triglyceride levels in the different genotypes (Fig. 3C). Thus, both loss-of-function and gain-of-function experiments indicate that miR-7 stimulates the storage of fats in growing larvae.

### miR-7 inhibits dILP2 synthesis and release

miR-7’s effects on fat stores, circulating sugars, and growth suggest that it might negatively regulate dILPs within IPCs. A genomic transgene containing the *dILP2* transcription unit and regulatory sequences is sufficient to functionally replace the loss of the endogenous *dILP2* gene^25^. Moreover, this transgene has been modified to place epitope tags fused to the dILP2 product. Like insulin, dILP2 is composed of A-chain and B-chain peptides linked via disulfide bonds. With a FLAG tag fused to the amino terminus of the A chain and an HA epitope fused to the carboxy terminus of the B chain, the resulting product is normally processed, secreted, and fully functional^25^. This *dILP2HF* transgene enabled us to monitor both mRNA and peptide expressed from the IPCs. First, we asked whether *dILP2HF* mRNA expression changed as a result of miR-7 perturbation. *dILP2HF* mRNA abundance was measured in whole brains of late third instar larvae by RT-qPCR. Sponge-mediated knockdown of miR-7 resulted in an increase in *dILP2HF* mRNA levels (Fig. 4A). Conversely, overexpression of miR-7 decreased *dILP2HF* mRNA abundance by three-fold (Fig. 4A). Thus, miR-7 represses *dILP2* mRNA levels.

**Figure 4.**
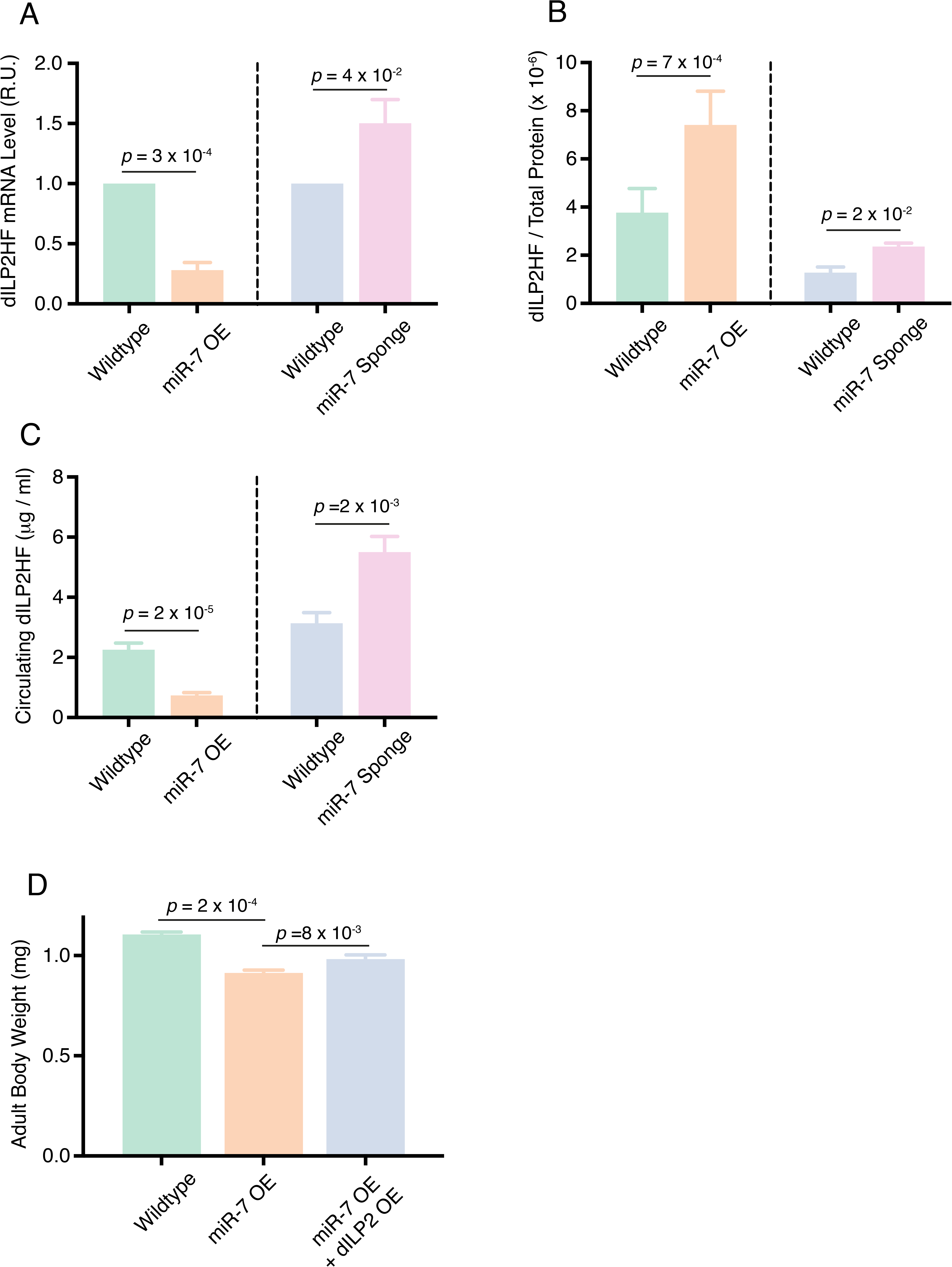
mir-7 mutants affect dILP2 expression and release. (**A**) Normalized levels of *dILP2HF* mRNA in larval brains from animals that have miR-7 overexpression (OE) or the miR-7 Sponge expressed in IPCs. Each condition is paired with a wildtype control line that contains the *dILP2-Gal4* driver used in each experiment. Measurements made by RT-qPCR are presented in relative units. (**B**) Fraction of total brain protein that is dILP2HF peptide (w/w) in larvae that have miR-7 overexpression (OE) or the miR-7 Sponge expressed in IPCs. Each condition is paired with a wildtype control line that contains the *dILP2-Gal4* driver used in each experiment. Different *dILP2-Gal4* drivers affect dILP2HF levels on their own, which requires rigorous pairing for control. (**C**) Concentration of dILP2HF peptide in larval hemolymph from animals that have miR-7 overexpression (OE) or the miR-7 Sponge expressed in IPCs. Each condition is paired with a wildtype control line that contains the *dILP2-Gal4* driver used in each experiment. For panels A - C, error bars represent the standard error of the mean, and *p* values are derived from unpaired two-tailed student T-tests. (**D**) Mean adult weight of females that have miR-7 overexpression (OE) compared to wildtype. One set of miR-7 OE animals also had dILP2 overexpressed in their IPCs. Error bars represent the standard error of the mean, and *p* values are derived from unpaired two-tailed student T-tests.

To determine if these effects were also exerted at the peptide level, we performed an ELISA assay, which can accurately measure dILP2HF peptide ranging from 40 attomoles to 4 femtomoles^25^. We observed a two-fold increase in dILP2HF peptide within the brain when miR-7 was knocked down by the sponge (Fig. 4B), consistent with effects on mRNA levels. Surprisingly, we saw that dILP2HF peptide level within the brain also increased when miR-7 was overexpressed in IPCs, even though mRNA levels decreased (Fig. 4A,B). Therefore, we measured the levels of dILP2HF peptide in circulation. Knocking down miR-7 in IPCs elevated circulating dILP2HF levels consistent with its effects on brain dILP2HF (Fig. 4C). Strikingly, miR-7 overexpression decreased the levels of circulating dILP2HF (Fig. 4C). Thus, when miR-7 was overexpressed in IPCs, stored brain dILP2HF rose but circulating levels dropped. This suggested that miR-7 might inhibit the release of dILP2HF from IPCs, thereby accounting for a buildup of stored dILP2HF and a drop of dILP2HF in circulation.

Our results suggest that miR-7 inhibits body growth, and miR-7 also inhibits dILP2 synthesis and release from IPCs. We wondered if the effect of miR-7 on dILP2 was responsible for its effect on growth. To test this idea, we overexpressed dILP2 in IPCs that also overexpressed miR-7 (Fig. 4D). The body weight of such individuals was partially rescued from the miR-7 effect. Thus, dILP2 at least partly mediates the effect of miR-7 on growth. One possible explanation for the partial rescue is that dILP2 overexpression does not necessarily alleviate the impact miR-7 has on dILP2 release from the IPCs.

### Identification of *cpa* as a miR-7 regulatory target

MicroRNAs repress their target genes by imperfect base-pairing to the 3’UTR of mRNAs. The primary recognition sequence is a 7 - 8 nucleotide "seed" that anneals to the 5’ end of a miRNA^48^. To identify mRNAs that are directly regulated by miR-7, we used computational prediction based on the presence of conserved seed sequences in annotated mRNAs. Different algorithms use different criteria when predicting the interaction between a miRNA and its mRNA target. To reduce the risk of identifying false positives, five different target prediction algorithms were used in parallel. We initially asked whether any algorithm predicted that miR-7 directly represses *dILP2* expression by base-pairing to *dILP2* mRNA. Our rationale was that since miR-7 inhibits *dILP2HF* mRNA expression, it might do so directly. However, none of the algorithms predicted miR-7 binding sites in the 3’UTR of the *dILP2, 3* and *5* genes (data not shown).

Predicted target genes were prioritized by considering those genes identified by three or more different algorithms. A total of 97 genes were identified that fulfilled this criterion. We next performed an experimental screen of these genes to determine if any of them might regulate IPC function. If a target gene directly mediates the effect of miR-7 on growth control by IPCs, we reasoned that RNAi knockdown of the gene specifically in IPCs would resemble miR-7 overexpression. The simplest phenotypic readout of growth control is adult body size, and miR-7 overexpression causes adults to weigh less than normal (Fig. 2A). Adult body size scales proportionally with wing blade length: the distance between points where the third longitudinal vein intersects the wing margin and anterior crossvein (Fig. 5A)^49^. miR-7 overexpression in IPCs reduced wing blade length as expected (Fig. 5B). We crossed *dILP2-Gal4* to *UAS-RNAi* lines that knocked down individual candidate miR-7 target genes, and measured blade length of affected animals. Of the RNAi lines tested, one resulted in a significant decrease in blade length comparable to miR-7 overexpression (Fig. 5B). The RNAi line knocked down *capping protein alpha* (*cpa*), a gene that encodes F-actin-capping protein subunit alpha. CPA binds the barbed ends of actin filaments to block their assembly and disassembly^50^. Unlike other capping proteins, CPA does not sever F actin. Although CPA is not known to regulate insulin, the actin cytoskeleton is a known regulator of insulin secretion, and disrupting other actin capping proteins has been shown to cause changes in insulin secretion^51,52^.

**Figure 5.**
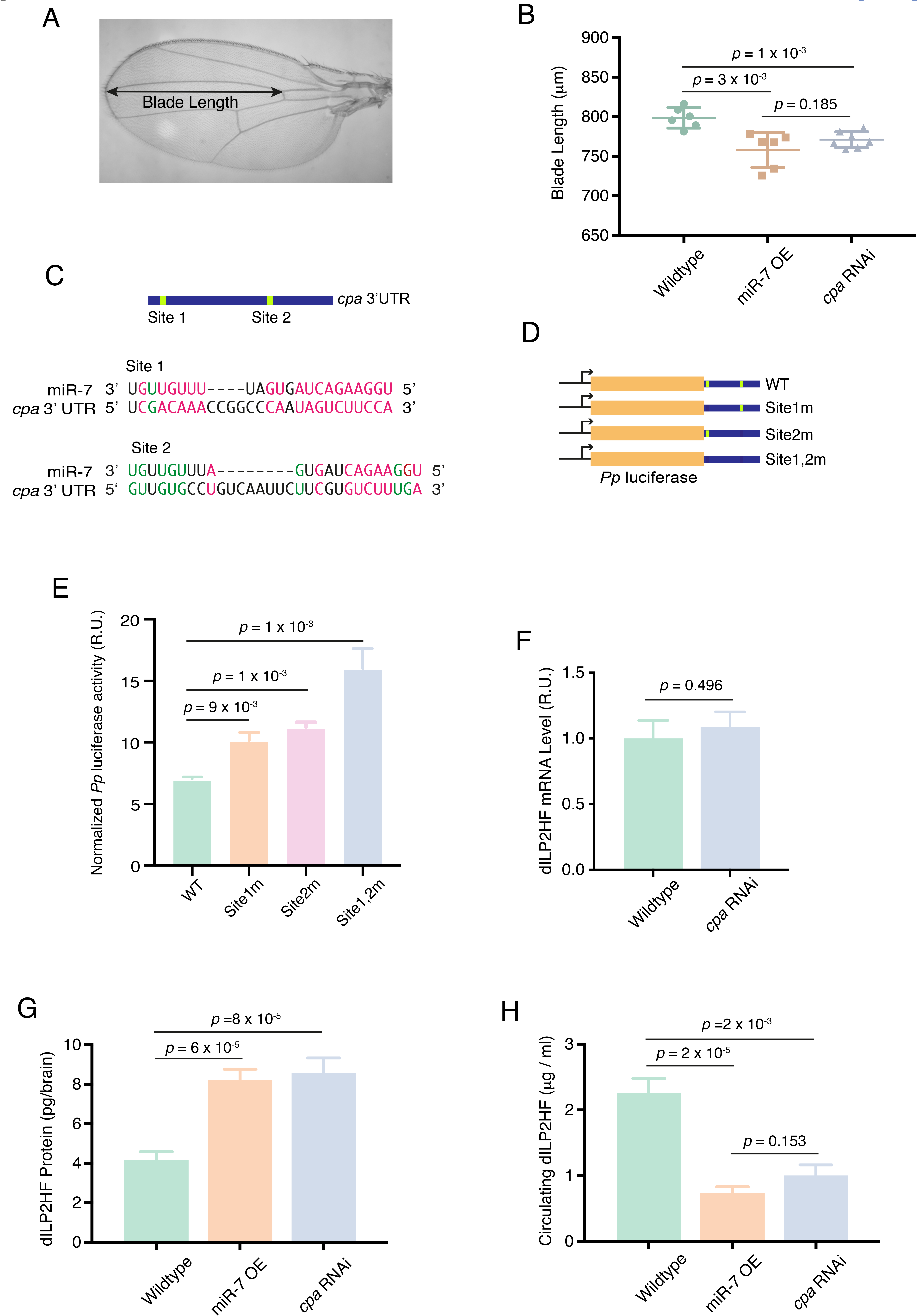
miR-7 regulates *cpa* expression to control IPC activity. (**A**) An adult wing showing the measurement of blade length, from the margin to the anterior crossvein where both intersect the L3 vein. (**B**) Blade lengths of females that contain *dILP2-Gal4* and either overexpress miR-7 (OE) or an RNAi hairpin directed against the *cpa* gene. Error bars are standard deviations, and *p* values are from two-tailed student T-tests. (**C**) Schematic showing the *cpa* 3’UTR, which contains two predicted miR-7 binding sites. Alignments show *Drosophila* miR-7-5p strand paired to binding sites 1 and 2 at nucleotide level resolution. Watson-Crick base-pairs are highlighted in red and GU wobble base-pairs are highlighted in green. **(D)** Schematic showing firefly luciferase fused to the *cpa* 3’UTR in reporter genes. Wildtype (WT) and mutant 3’UTRs used in reporters are shown. (**E**) Firefly luciferase activity from the different reporter genes normalized to Renilla luciferase in S2 cells expressing miR-7. (**F**) *dILP2HF* mRNA levels in larval brains from wildtype animals or animals that have a RNAi knockdown of *cpa* in IPCs. Measurements made by RT-qPCR are presented in relative units. (**G**) Stored brain dILP2HF peptide (by weight) in larvae that have miR-7 overexpression (OE) or RNAi knockdown of *cpa* in IPCs. There was no significant difference between miR-7 OE and *cpa* RNAi levels of dILP2HF (*p* = 0.76, t test). (**H**) Concentration of dILP2HF peptide in larval hemolymph from animals that have miR-7 overexpression (OE) or RNAi knockdown of *cpa* in IPCs. For panels E - H, error bars represent the standard error of the mean, and *p* values are derived from unpaired two-tailed student T-tests.

There are two predicted miR-7 binding sites in the *cpa* 3’UTR. Site 1 has an 8mer seed sequence and shows strong potential binding to miR-7 (Fig. 5C). Site 2 has an offset 6mer seed sequence, which is not predicted to bind as strongly, though the site has 3’ compensatory pairing (Fig. 5C). To verify that *cpa* is a direct target of miR-7, we fused the *cpa* 3’UTR to a firefly luciferase reporter gene (Fig. 5D). We also generated reporters with base mutations in the seeds of site 1 only, site 2 only, or both sites. We then transfected these reporters into *Drosophila* S2 cells, along with a Renilla luciferase control reporter, and miR-7 expression constructs. Mutation of either site 1 or site 2 resulted in a 1.5 fold derepression of firefly luciferase expression (Fig. 5E). Mutation of both sites resulted in greater derepression, suggesting that the sites function additively to inhibit gene expression (Fig. 5E). Taken together, these results demonstrate that the sites can directly mediate miR-7 regulation of *cpa* transcripts.

If miR-7 acts through *cpa* to regulate IPC function, we predicted that RNAi of *cpa* should phenocopy the effects seen with miR-7 overexpression in IPCs. Therefore, we tested *cpa* RNAi animals for the other phenotypes observed with miR-7 overexpression. First, we quantified *dILP2* transcript levels in IPCs of RNAi animals and found that the *cpa* RNAi line showed no change in *dILP2* mRNA levels (Fig. 5F). This result suggests that CPA is not involved in production of *dILP2* mRNA. However, when dILP2HF peptide levels were quantified in the brains of *cpa* RNAi animals, there was an increase in stored brain dILP2HF that was indistinguishable from miR-7 overexpression (Fig. 5G). Measurement of circulating dILP2HF showed that <u>*cpa*</u> RNAi decreased circulating dILP2HF to a degree indistinguishable from miR-7 overexpression (Fig. 5H). From these experiments we conclude that CPA is required to stimulate the release of dILP2 from IPCs.

### miR-7 regulation of CPA expression is conserved in mammalian β-cells

The mature miR-7 RNA has perfectly identical sequence conservation between *Drosophila* and mammals (Fig. 1A). Our findings indicate that miR-7 regulates dILP2 release in *Drosophila* IPCs, just as miR-7 regulates insulin secretion in mouse β-cells^28^. This led us to question whether the miR-7 regulation of *capping protein alpha* expression is similarly conserved in mammals. TargetScan, Pictar, and PITA were used to predict miR-7 target genes in the human genome. Using the same search parameters and stringency as the *Drosophila* search, we found 571 human genes that were predicted targets. We then determined whether each *Drosophila* and human candidate gene had an ortholog in the other species, and cross-referenced the two lists to find orthologs in both species that are predicted miR-7 targets. Of the 571 candidates in humans and 97 candidates in *Drosophila*, only 10 pairs of candidates were orthologous to one another. Interestingly, *cpa* and its human ortholog *CAPZA1* were the top candidates, in that all algorithms predicted *cpa* and *CAPZA1* to be miR-7 targets. There is one predicted miR-7 binding site in human *CAPZA1* (Fig. 6A). The mouse *Capza1* gene also has a predicted miR-7 binding site (Fig. 6A).

**Figure 6.**
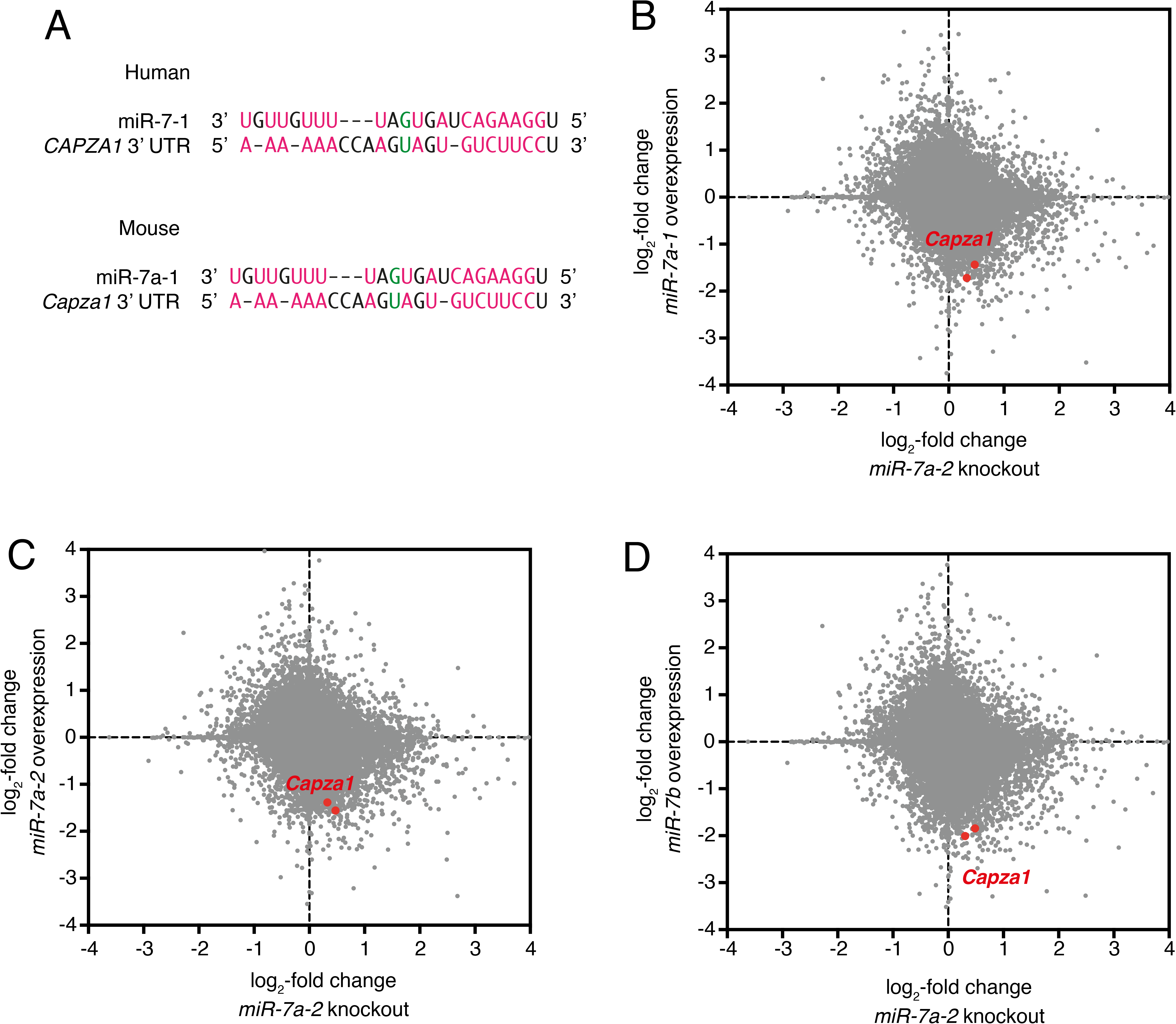
miR-7 regulates *Capza1* expression in mouse pancreatic β-cells. (**A**) Alignment of the human miR-7-1-5p strand with its predicted binding site in the 3’ UTR of human *CAPZA1*. Alignment of the mouse miR-7a-1-5p strand with its predicted binding site in the 3’ UTR of mouse *Capza1*. Watson-Crick base-pairs are highlighted in red and GU wobble base-pairs are highlighted in green. (B-D) Scatter plots of all detected mRNAs (grey) in samples of mouse β-cells. Data is from Latreille et al., (2014)^28^. The Y axis of each plot shows the log_2_fold difference in mRNA levels from MIN6 cell lines overexpressing a miR-7 paralog vs. not overexpressing the paralog. The X axis of each plot shows the log_2_fold difference in mRNA level pancreatic islets isolated from *miR-7a-2* knockout mice vs. wildtype mice. Highlighted in red are the two replicate samples measured for the *Capza1* mRNA in each experiment. (**B**) MIN cells overexpressing miR-7a-1 vs. miR-7a-2 knockout islets. **(C)** MIN cells overexpressing miR-7a-2 vs. miR-7a-2 knockout islets. **(D)** MIN cells overexpressing miR-7b vs. miR-7a-2 knockout islets.

To determine whether miR-7 regulates *capping protein alpha* expression in mammalian pancreatic β-cells, we turned to data from mouse experiments^28^. Both humans and mice contain three paralogous genes encoding miR-7 isoforms. The major miR-7 paralogs are identical in sequence between the two species (Fig. 1A), and they are predicted regulators of *capping protein alpha* homologs in both mouse (*Capza1*) and human (*CAPZA1*) (Fig. 6A). Therefore, we analyzed a published dataset of genome-wide transcript levels from mouse β-cells^28^. This dataset contained transcriptomics from wildtype β-cells, as well as β-cells in which *mir-7* paralogs were either knocked out or overexpressed. We focused our attention on *Capza1* mRNA levels that were quantitated in these samples. Strikingly, there was an increase in *Capza1* mRNA levels in *mir-7a-2* knockout β-cells (Fig. 6B-6D). Conversely, there was a decrease in *Capza1* mRNA levels when any one of the three miR-7 paralogs was overexpressed in β-cells (Fig. 6B-6D). This result indicates that mouse *Capza1* is repressed by miR-7 in pancreatic β-cells.

## Discussion

Of the thousands of miRNAs in the human genome, very few show as high a degree of evolutionary sequence conservation as miR-7^33^. This has led us to question whether miR-7’s sequence conservation is coupled to a conserved biological function. In mouse β-cells, miR-7 inhibits glucose-stimulated insulin secretion^28^. We have found that in *Drosophila*, miR-7 inhibits dILP2 peptide release from IPCs. Thus, miR-7 plays a highly conserved role in negatively regulating insulin secretion. *Drosophila* miR-7 appears to do so by repression of CPA protein expression gene in IPCs. We have found that knockdown of CPA in *Drosophila* IPCs interferes with secretion of dILP2 peptide into hemolymph. CPA binds to the barbed ends of actin filaments to inhibit additional polymerization or depolymerization^50^. Dynamic actin filament polymerization is known to be important for proper insulin secretion^51,52^. Mouse cell lines that have a mutation in the actin-binding protein gelsolin are unable to undergo glucose-mediated insulin exocytosis due to their inability to depolymerize their cytoskeleton^51^. It remains unclear exactly how loss of CPA disrupts the secretory process, and whether it occurs via actin capping. Perhaps CPA stimulates dILP2 secretion by similar mechanisms, and miR-7 attenuates CPA abundance, thereby down-regulating dILP2 secretion.

This then raises the question as to whether miR-7 attenuation of mouse *Capza1* gene expression also downregulates insulin secretion in pancreatic β-cells. Although SNCA was found to mediate the effects of mouse miR-7a-2 on insulin secretion^28^, it appears that mouse miR-7a-2 also directly represses *Capza1* expression. If Capza1 plays a role in insulin secretion, then the conserved regulatory interaction between miR-7 and CPA/Capza1 is likely coupled to the conserved endocrine control of glucose metabolism.

In the marine annelid *Platynereis*, miR-7 is expressed in neurosecretory cells that release the vasotocin and FMRF-amide neuropeptides^53^. Although FMRF-amides are exclusive to invertebrates, vasotocin is homologous to oxytocin and vasopressin in mammals, and vasotocin is secreted from the pituitary gland in non-mammalian vertebrates. Strikingly, the regions of the vertebrate brain that secrete these neuropeptides, the hypothalamus and pituitary, are also regions where the miR-7 gene family is highly expressed^54^. miR-7a2 was recently shown to regulate hormone secretion in the mouse pituitary^55^. Possibly, an ancient role for miR-7 was to regulate neuropeptide release in the bilaterian ancestor, and this role has been retained during the evolution of all animals. Interestingly, the actin binding protein profilin seems to be regulated by miR-7 in both pancreatic islet cells and in the pituitary^55^ suggesting that perhaps miR-7 has retained a role in regulating the cytoskeleton across organisms and in different endocrine cell types.

## Acknowledgements

We thank the Seung Kim Lab for generously providing us with the protocol, the reagents and the *Drosophila* stock to quantify stored and circulating dILP2HF. We additionally thank members of the Carthew and Beitel Labs for helpful discussions and advice. P.A. and J.J.C. were supported in part by the Cell and Molecular Basis of Disease (CMBD) Training Grant (T32GM008061). R.W.C. was supported by R35GM118144.

